# Automatic Detection of Synaptic Partners in a Whole-Brain *Drosophila* EM Dataset

**DOI:** 10.1101/2019.12.12.874172

**Authors:** Julia Buhmann, Arlo Sheridan, Stephan Gerhard, Renate Krause, Tri Nguyen, Larissa Heinrich, Philipp Schlegel, Wei-Chung Allen Lee, Rachel Wilson, Stephan Saalfeld, Gregory Jefferis, Davi Bock, Srinivas Turaga, Matthew Cook, Jan Funke

## Abstract

The study of neural circuits requires the reconstruction of neurons and the identification of synaptic connections between them. To scale the reconstruction to the size of whole-brain datasets, semi-automatic methods are needed to solve those tasks. Here, we present an automatic method for synaptic partner identification in insect brains, which uses convolutional neural networks to identify post-synaptic sites and their pre-synaptic partners. The networks can be trained from human generated point annotations alone and require only simple post-processing to obtain final predictions. We used our method to extract 244 million putative synaptic partners in the fifty-teravoxel full adult fly brain (FAFB) electron microscopy (EM) dataset and evaluated its accuracy on 146,643 synapses from 702 neurons with a total cable length of 312 mm in four different brain regions. The predicted synaptic connections can be used together with a neuron segmentation to infer a connectivity graph with high accuracy: between 92% and 96% of edges linking connected neurons are correctly classified as weakly connected (less than five synapses) and strongly connected (at least five synapses). Our synaptic partner predictions for the FAFB dataset are publicly available, together with a query library allowing automatic retrieval of up- and downstream neurons.

## 1 Introduction

Obtaining a connectome requires the reconstruction of neurons and the identification of synaptic connections between them. Currently, high-resolution electron microscopy (EM) is the only scalable method to reliably resolve individual synaptic connections. As shown in Fig. 2a, EM at nanometer scale provides enough detail to identify pre-synaptic sites, synaptic clefts, and post-synaptic partners.

**Figure 1:**
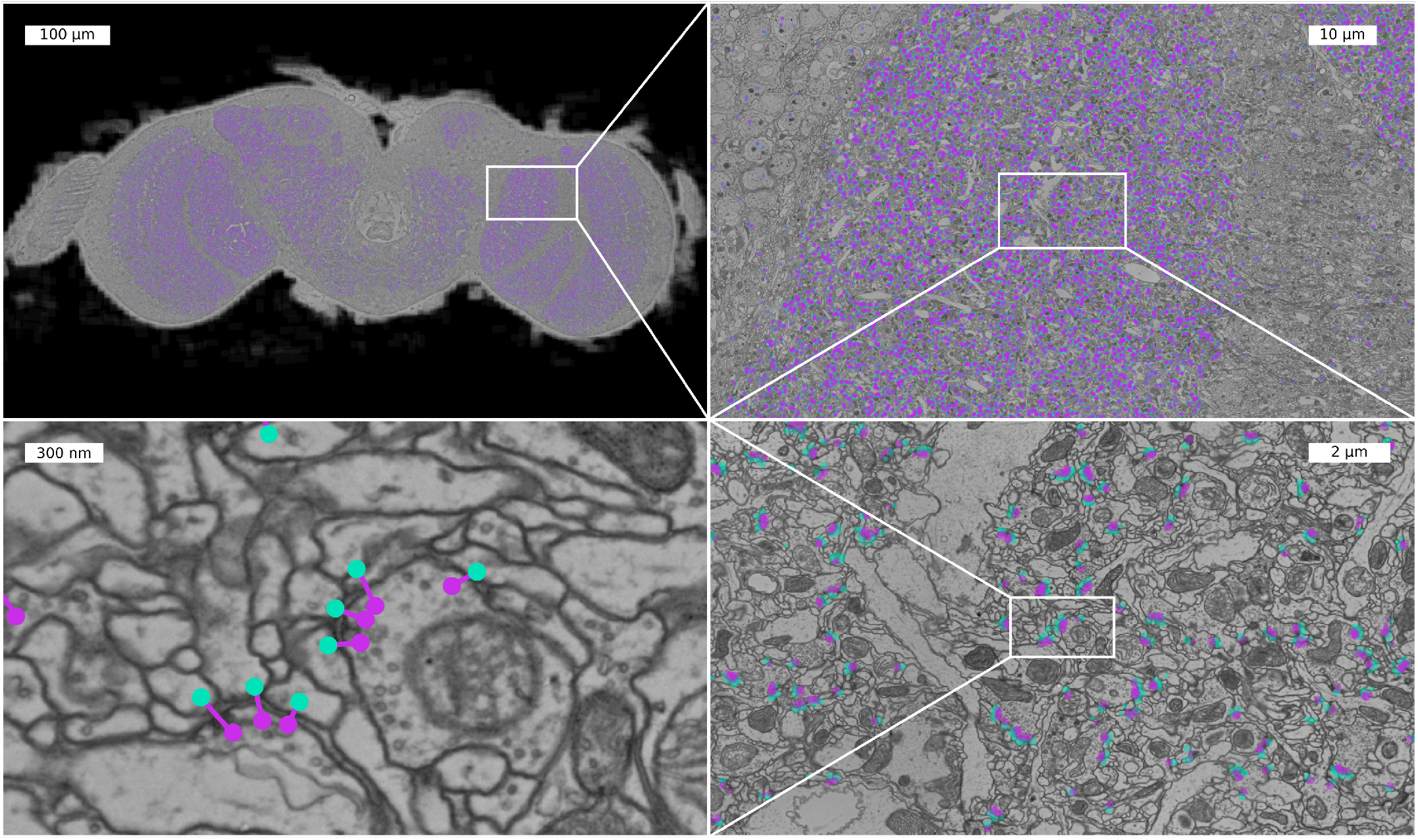
Sample section of the FAFB dataset with predicted synaptic partners. Sequence of increasing zoom levels of a coronal section (z-section, sectioning plane) with predicted connections (pre-synaptic site purple, post-synaptic site turquoise).

**Figure 2:**
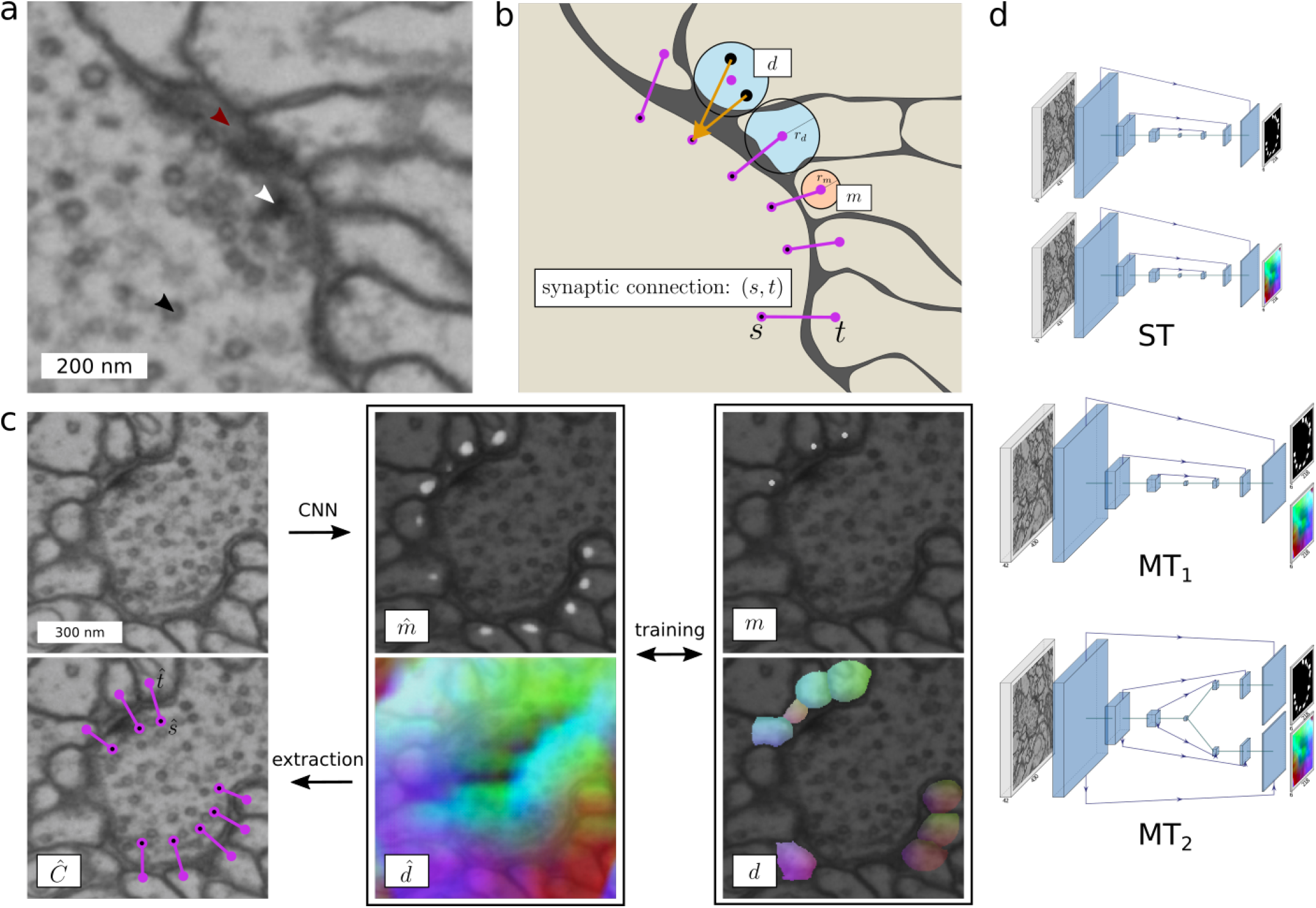
Synaptic partner representation and corresponding CNN architectures. **a**) Appearance of a polyadic synapse in EM data with vesicles (black arrow), T-Bar (white arrow), and synaptic cleft (red arrow). **b**) Given ground-truth annotations (*s*, *t*) for synaptic partner sites, we generate a post-synaptic site mask *m* (indicating the location of post-synaptic sites) and a field of 3D vectors *d* (pointing to the corresponding pre-synaptic site). Created spheres around point annotations shown in peach for *m* and in light blue for *d*. **c**) We train a CNN on *m* and *d* to predict 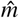 and 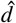, from which we extract synaptic connections. 3D vectors in *d* and 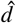 are RGB-color encoded. **d**) Investigated CNNs: ST (one U-Net per 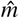 and 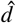), MT_1_ (one U-Net for both 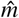 and 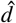), and MT_2_ (one U-Net with two separate upsampling paths per 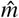 and 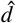).

Connectome analysis of the fruit fly is of particular interest on a whole-brain scale. So far, the FAFB (full adult fly brain) EM volume (Zheng et al., 2018) is the only available dataset comprising a complete adult fly brain. Reconstruction of neuron morphologies and synaptic connections in this volume has been done manually so far, totalling at the time of writing ~1.2 million neurons and neuron fragments with a total length of ~10.7 m and ~1.1 million synaptic connections, traced out in a collaborative effort over a period of four years. Clearly, the manual analysis of EM volumes is the main bottleneck in the acquisition of connectomes, dwarfing even the significant time needed for sample preparation, imaging, and alignment in recent imaging pipelines (Graham et al., 2019).

Consequently, machine learning based methods have been developed to automate the analysis of EM volumes. Over the last decade, the focus of these efforts has mainly been the segmentation of neurons, which are arguably the more time consuming structures to reconstruct (for an overview of recent methods see the reviews by Lee et al. (2019) and Sheridan et al. (2019)). Tracing, for example, a Kenyon cell by hand in FAFB inside the calyx takes about 46.7 s/μm on average (Li et al., 2019), resulting in a total of 1094 days for all of the approximately 5000 Kenyon cells neurons in an adult fly brain (Fahrbach, 2006). Identification of synaptic connections, on the other hand, requires about 5 s per connection, resulting in a total of 32 days for all of the approximately 570,000 incoming Kenyon cell connections. Recent neuron segmentation algorithms provide initial segmentations that significantly reduce the time needed to trace a Kenyon cell to about 8.6 s/μm (Li et al., 2019) or 201 days for all Kenyon cells. The relative time spent on manually identifying synaptic connections has thus increased significantly (Huang et al., 2018; Dorkenwald et al., 2017), highlighting the need for automatic methods for synapse annotation.

To meet this need, several automatic approaches have been proposed, tailored towards the model organism of interest. In vertebrates, recent methods have been shown to be accurate enough to directly infer synaptic connectivity reliably (Dorkenwald et al., 2017; Motta et al., 2019). In insect brains, however, the task of identifying synaptic connections is more challenging, since the sizes of synapses is generally smaller than in vertebrates and many synapses are *polyadic* (*i.e.*, one pre-synaptic site connects to several post-synaptic sites). Most recent methods for synaptic partner prediction deal with those challenges in two steps: The first step consists of identifying and localizing synaptic features such as T-bars or synaptic clefts (visible in Fig. 2a), whereas in the second step neuron segments (as induced by a neuron segmentation) in physical contact are classified as synaptically “connected” or “not-connected”. For the first step, convolutional neural networks (CNNs) were successfully used to detect synaptic clefts (Heinrich et al., 2018) or T-bars (Huang et al., 2018) in large-scale fruit fly EM data. For the second step, Kreshuk et al. (2015) proposed a graphical model to solve synaptic partner assignment given a neuron segmentation and candidate synapse detections. More recently, CNNs have been used to infer synaptic connectivity: Turner et al. (2019) use a CNN to predict pre- and post-synaptic neuron masks from synaptic cleft segmentations and Parag et al. (2018) use a CNN to classify neuron pair candidates extracted from a neuron segmentation as “connected” or “not-connected”. Huang et al. (2018) solve a similar classification task with a multilayer perceptron and local context features. In contrast to these two-step approaches, Buhmann et al. (2018) jointly detect pre- and post-synaptic sites and infer their connectivity via long-range affinity edges using a single CNN architecture.

However, few methods have been applied so far to large-scale *Drosophila* datasets: Heinrich et al. (2018) densely predict synaptic clefts in the whole FAFB dataset, albeit without extraction of synaptic partners. To the best of our knowledge, only the method proposed by Huang et al. (2018) has been used to predict synaptic partners in a large *Drosophila* dataset with a size of 128k μm^3^.

Here, we present a novel method for synaptic partner identification, which can be trained from human generated point annotations alone. More specifically, we use a 3D U-Net CNN (Ronneberger et al., 2015) to predict for each voxel in a volume whether it is part of a post-synaptic site and, if it is, an offset vector pointing from this voxel to the corresponding pre-synaptic site. The method we introduce here is similar to our previous work (Buhmann et al., 2018) in that it does not require explicit annotation or detection of synapse features (such as T-bars or clefts), but instead predicts synaptic partners from raw EM data directly. This design choice has two practically important implications: First, our method can be trained from pre- and post-synaptic point annotations alone, which reduces the effort needed to annotate future training datasets. Second, the prediction of synaptic partner candidates does not rely on the availability of a neuron segmentation, which makes our method complementary to, but not dependent on, a high-quality neuron segmentation.

We used our method to extract 244 million putative synaptic partners in the complete FAFB dataset and evaluated its accuracy on 146,643 synapses from 702 neurons with a total cable length of 312 mm in four different brain regions. When measuring performance at an individual synaptic connection level, we obtain f-scores of 0.73, 0.68, 0.66 and 0.59 for the four different brain areas. We observe a decrease in f-score in regions farther away from the training datasets, suggesting that more diverse training datasets are needed to improve overall accuracy. When performance is measured in the context of connectome inference, we find that our predicted synaptic connections can be used to automatically infer a connectivity graph with high accuracy: 96% of edges between connected neurons are correctly classified as strong (five synaptic connections or more) vs. weak (less than five synaptic connections). Source code, predictions, and evaluation datasets are publicly available^1^. To facilitate circuit reconstruction efforts, we additionally provide a query library that allows automatic inference of up- and down-stream neurons based on our predictions as part of FafbSeg-Py^2^.

## 2 Method

### 2.1 Synapse Representation

Due to the polyadic nature of synapses in *Drosophila*, we represent synapses as sets of connections *C* ⊂ Ω^2^ between pairs of voxels (*s*, *t*) ∈ Ω^2^, where *s* and *t* denote pre- and post-synaptic locations, respectively, and Ω the set of all voxels in a volume. In this representation, pre-synaptic locations do not have to be shared for connections that involve the same T-bar or synaptic cleft (Fig. 4a). This is in constrast to the convention followed in annotation tools like Catmaid or NeuTu, where one pre-synaptic connector node serves as a hub for all post-synaptic partners (Fig. 4e). Although this convention might be beneficial for manual annotation, the representation proposed here is potentially better suited for automatic methods, as it does not require the localization of a unique pre-synaptic site per synapse. Instead, automatic methods can learn to identify synaptic partners independently from each other. Furthermore, this representation is also the one used by the Cremi challenge, which allows us to use the provided training data and evaluation metrics.

### 2.2 Connection Prediction

Given ground-truth annotations *C* and raw intensity values *x* : Ω ↦ ℝ, we train a neural network to produce two voxel-wise outputs: a mask 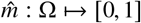 to indicate post-synaptic sites and a field of 3D vectors 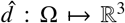 that point from post-synaptic sites to their corresponding pre-synaptic site (see Fig. 2 for an illustration). Both outputs are trained on ground-truth *m* and *d*, extracted from *C* in the following way: *m*(*i*) is equal to 1 if voxel *i* is within a threshold distance *r_m_* to any post-synaptic site *t* over all pairs (*s*, *t*) ∈ *C*, and 0 otherwise (resulting in a sphere). Similarly, *d*(*i*) = *s* − *i* for every voxel *i* within a threshold distance *r_d_* to the closest *t* over all pairs (*s*, *t*) ∈ *C*, and left undefined otherwise (during training, gradients on 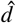 are only evaluated where *d* is defined). To obtain a set 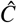 of putative synaptic connections from predictions 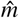 and 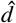, we find connected components in 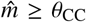. For each component, we sum up the underlying predicted values of 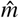, which we refer to as the *connection score*. Each component with a score larger than a threshold *θ*_CS_ is considered to represent a post-synaptic site, from which we extract a location 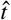 as the maximum of an Euclidean distance transform of the component. The corresponding pre-synaptic site location 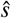 is then found by querying the predicted direction vectors 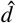, *i.e.*, 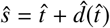.

### 2.3 Network Architectures

We use a 3D U-Net (Falk et al., 2019) as our core architecture, following the design used by Funke et al. (2018) and Heinrich et al. (2018), *i.e.*, we use four resolution levels with downsample factors in *xyz* of (3, 3, 1), (3, 3, 1), and (3, 3, 3) and a five-fold feature map increase between levels. Convolutional passes are comprised of two convolutions with kernel sizes of (3, 3, 3) followed by a ReLU activation. Final activation functions for 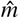 and 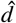 are sigmoid and linear, respectively. The initial number of feature maps *f* is left as a free hyper-parameter. In order to train the two outputs 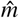 and 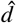 required by our method, we explore three specific architectures: Architecture ST (single-task) refers to two separate U-Nets trained individually for either 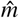 and 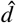, and thus does not benefit from weight sharing. Architectures MT_1_ and MT_2_ refer to multi-task networks that use weight sharing to jointly predict 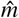 and 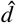. MT_1_ is a default U-Net with four output feature maps (one feature map for 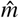 and three feature maps for 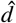). MT_2_ is a U-Net with two independent upsampling paths (as independently proposed by Wang et al. (2019)), each one predicting either 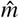 and 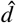, *i.e.*, weights are only shared during the downsampling path between the two outputs. Fig. 2d summarizes the architecture choices.

### 2.4 Training

Our training loss *L_d_* for (*d*, 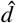) is based on mean squared error (MSE). For the post-synaptic mask loss 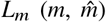, we explore both error functions MSE and cross-entropy (CE). In the multi-task context (MT_1_, MT_2_), we simply sum the two losses to obtain the total loss: *L_total_* = *L_d_* + *L_m_*.

We train all investigated networks (ST, MT_1_, and MT_2_) on randomly sampled volumes of size (430, 430, 42) voxels, which we refer to as *mini-batches*. Each mini-batch is randomly augmented with xy-transpositions, xy-flips, continuous rotations around the z-axis, and section-wise elastic deformations and intensity changes. We use the Gunpowder library^3^ for data loading, preprocessing, and all augmentations mentioned above. Training using Tensor-Flow on an NVIDIA Titan X GPU takes 6-11 days to reach 700,000 iterations.

Due to the sparseness of post-synaptic sites, the training signal *m* is heavily unbalanced: ~65% of randomly sampled mini-batches do not contain post-synaptic sites at all, the remaining mini-batches have a ratio of foreground- to background-voxels of ~1:4000 on average. We mitigate this imbalance with two balancing strategies: First, we reject empty mini-batches during training with probability *p*_rej_. Second, we scale the loss for 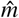 voxel-wise with the inverse class frequency ratio of foreground- and background-voxels, clipping at 0.0007^−1^. In practice, the loss for a foreground voxel is scaled on average with a factor of ~1400.

## 3 Results

We use the proposed method to process the entire FAFB dataset. The FAFB dataset contains a complete female *Drosophila* brain, recorded at a resolution of (4, 4, 40) nm, imaged using serial section transmission electron microscopy (Zheng et al., 2018). The volume’s dimensions, including background, are (992, 535, 282) μm, although only ~20% of its volume contains neural tissue (Li et al., 2019). See (Zheng et al., 2018) for more details about the FAFB dataset.

The following section provides details about the training and validation procedure for model selection, our pipeline for processing large volumes, and quantitative results on five evaluation datasets spanning four different brain regions.

### 3.1 Model Validation

We train and validate our method using the publicly available training dataset of the Cremi challenge^4^. This dataset consists of three 5μm cubes, cropped from the calyx (part of the mushroom body) of the FAFB volume, in which synaptic partners were fully annotated (1965 synapses in total, see Fig. 4a for examples). We realign and split those cubes into a training dataset (75% of the volume of each cube, resulting in a total of 1402 synapses) and a validation dataset (25%, 540 synapses). As Cremi also provides neuron segmentation, we intersect the spheres around point annotations in *m* and *d* with the volume of the respective neuron, although this step is not necessary because we obtain similar performance when using no neuron segmentation for training (data not shown).

We use the Cremi evaluation procedure to assess performance. In short, we match a predicted connection to a ground-truth connection if the underlying neuron segmentation IDs are the same and if the synaptic sites are in close distance (smaller than 400 nm). Matched connections are counted as true positives, unmatched ground-truth connections are considered false negatives, unmatched predicted connections are considered false positives. We obtain precision and recall values by adding up true positives, false positives, and false negatives of all three Cremi samples before computing precision and recall values. This is different from the original proposed Cremi score, where precision and recall is computed for each sample individually and then averaged, which effectively weights samples based on the inverse of the number of synaptic connections they contain.

We performed a grid search on the validation dataset to find the best performing configuration amongst different architecture choices (ST, MT_1_, and MT_2_, see Section 2.3), number of initial feature maps (small: *f* = 4, big: *f* = 12), training loss for 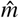 (CE: cross entropy, MSE: mean squared error), and two sample balancing strategies (standard: *p*_rej_ = 0.95 throughout training, curriculum: *p*_rej_ = 0.95 until 90k training iterations, then *p*_rej_ = 0, see Section 2.4). The resulting numbers of trainable parameters for the investigated small networks are 22 M (ST), 11 M (MT_1_), 12 M (MT_2_) and for the big networks 192 M (ST), 96 M (MT_1_), 115 M (MT_2_).

Fig. 3 (left panels, “without neuron segmentation”) summarizes the grid search results in terms of best f-score obtained over all connection thresholds *θ*_CS_. Best validation results are obtained using big networks with curriculum learning (f-score 0.74, precision 0.72, recall 0.77). Differences between training losses (CE and MSE) as well as architectures ST and MT_2_ are marginal, however, architecture MT_1_ consistently failed to jointly predict 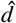 and 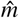 (results not shown), also when trained with a product loss (*L_total_* = *L_d_* · *L_m_*).

**Figure 3:**
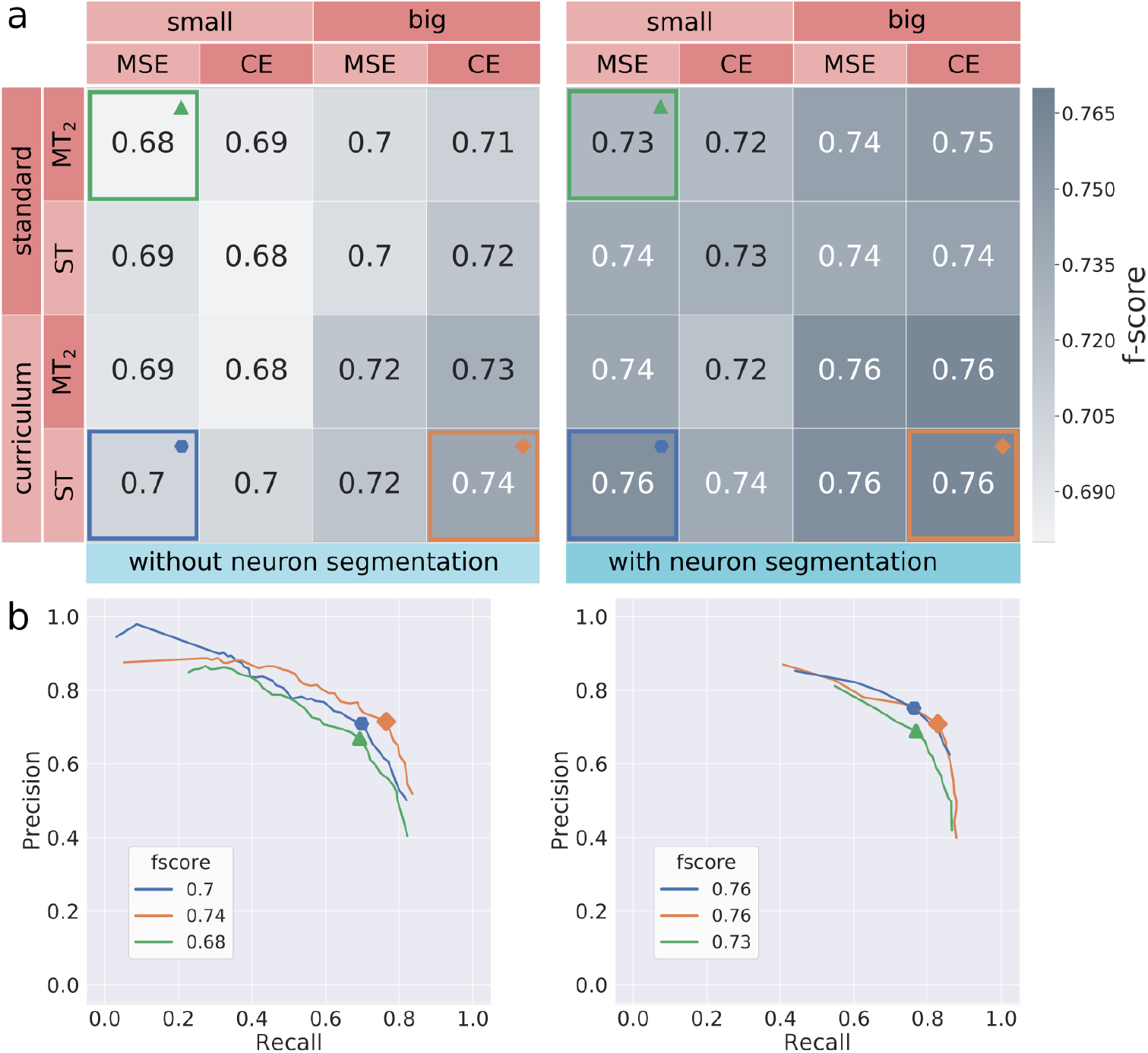
Validation results on Cremi dataset. **a**) Grid-search results in terms of best f-score (higher/darker is better) over all extraction thresholds for various parameter configurations: network size (small: *f* = 4, big: *f* = 12), training loss for 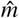 (MSE: mean squared error, CE: cross-entropy), balancing strategies (curriculum: *p*_rej_ = 0.95 until 90k training iterations then *p*_rej_ = 0, standard: *p*_rej_ = 0.95 constant), U-Net architectures (ST, MT_2_). Left side shows results without post-processing using a neuron segmentation. The best performing model (highlighted in orange) achieves an f-score of 0.74. Using a neuron segmentation for post-processing (right side), the best performing method improves to an f-score of 0.76 (orange highlight). Remarkably, previously inferior configurations are on par with the best performing one after post-processing (blue). **b**) Precision-recall curves over *θ*_CS_ for three selected models in (a), respective best f-score for each curve is highlighted.

**Figure 4:**
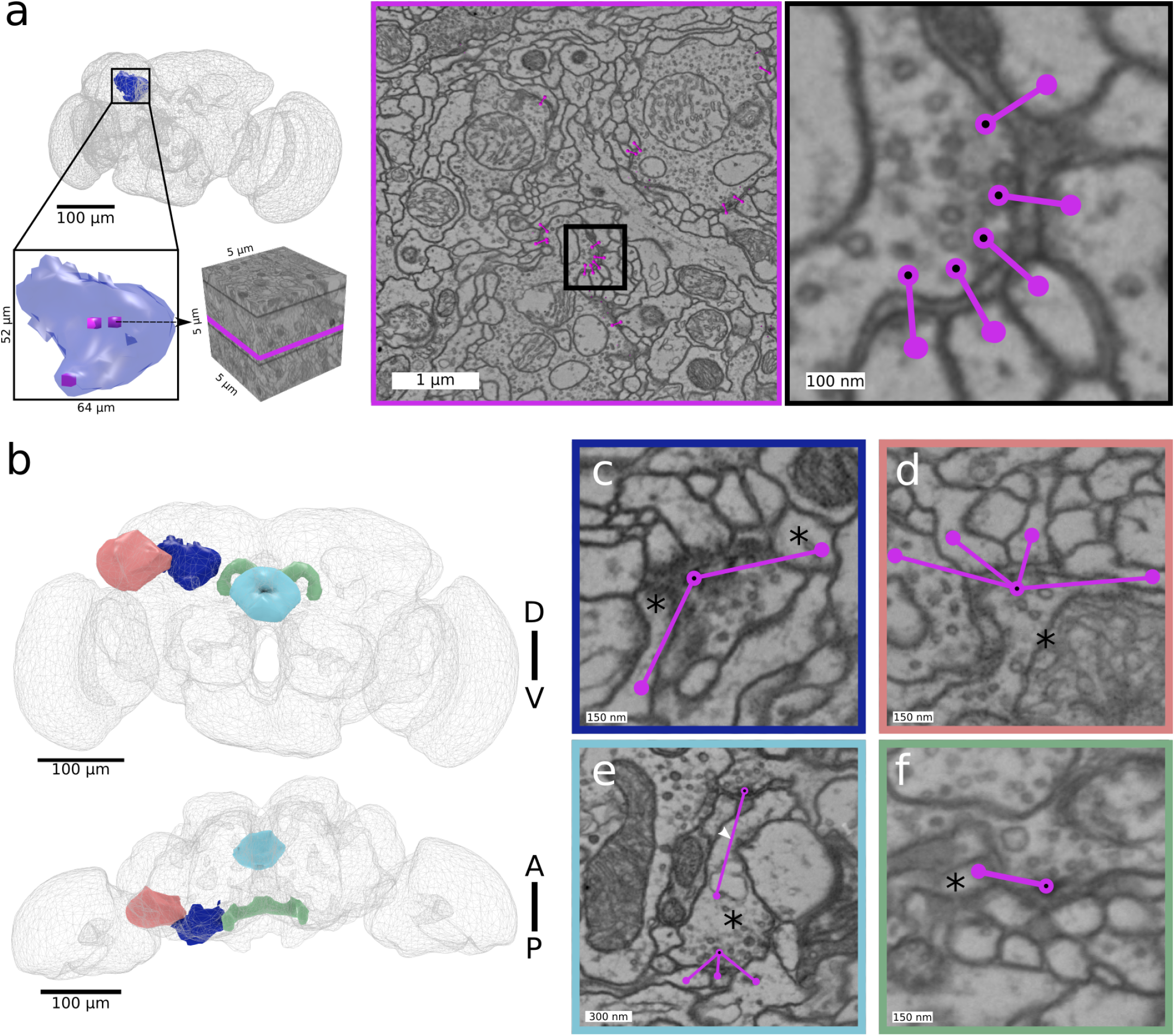
Synapse ground-truth datasets in FAFB used for training and validation (a) and evaluation (b-f). **a**) Three densely annotated ground-truth cubes (5×5×5 μm) located in the calyx brain area were used for training and model validation (Cremi training dataset). In this dataset, synaptic partners are annotated as individual pairs of pre- and post-synaptic sites (purple nodes, pre-synaptic sites marked with a black dot), connected by an edge. **b**) Four sparsely annotated ground-truth datasets located in the lateral horn (peach), calyx (dark blue), ellipsoid body (light blue), protocerebral bridge (green) were used for evaluation. **c**-**f**) Example synapse annotations for each of the four datasets from (b). Annotations are sparse and only complete for specific neurons (marked with a star). Pre- and post-synaptic locations are more distant than in (a). **c**) Incoming synapses of a Kenyon cell in calyx. **d**) Outgoing synapses of a projection neuron in lateral horn. **e**) Incoming- and outgoing synapses of a neuron in ellipsoid body. **f**) Incoming synapse of a neuron in protocerebral bridge. Rendering in a,b was done with with Catmaid-to-blender (Schlegel et al., 2016)).

### 3.2 Neuron segmentation improves performance

If a neuron segmentation is available for post-processing, as it is the case for the Cremi dataset, it can be used to improve accuracy of synaptic partner detection by filtering two types of false positives during post-processing: (1) false positives connecting the same neurite, and (2) duplicate close-by detections of a single synaptic partner pair across the same cleft.

The ability to remove duplicate detections around a cleft allows using local non-max suppression (NMS) to identify post-synaptic sites, which naturally gives rise to multiple detections in larger post-synaptic neurites. In addition to the hyperparameters discussed in the previous section, we therefore also test the impact of using NMS (radius: 40, 80, or 120 nm) for post-synaptic site detection compared to finding connected components (CC) as described in Section 2.2.

The benefits of using a segmentation for post-processing on the Cremi validation dataset are summarized in Fig. 3 (right panels, “with neuron segmentation”). Overall, scores improve (best f-score 0.76 compared to best f-score without neuron segmentation 0.74). Remarkably, smaller models (~90% less parameters) improve considerably to the point of matching the best observed score (small ST with curriculum learning using MSE improves f-score from 0.7 to 0.76). For each investigated combination of hyperparameters, NMS performed better than CC.

These findings suggest that in the presence of an accurate neuron segmentation smaller, more efficient architectures are on par with larger ones. It should be noted, however, that the neuron segmentation used here for validation was considerably proof-read and that the gain from purely automatic segmentations is likely less.

### 3.3 Synapse prediction on the complete FAFB dataset

Following the insights gained from validation on the Cremi datasets, we chose a small ST architecture with curriculum learning and a cross-entropy loss to process the whole FAFB volume. To limit prediction and synaptic partner extraction to areas containing synapses we ignore regions not part of the neuropil using the mask from Heinrich et al. (2018). The remaining volume consists of ~50 teravoxels or equivalently ~32 million μm^3^, which we process in blocks. For each block, we read in raw data with size (1132, 1132, 90) voxels to produce predictions of size (864, 864, 48) voxels (size difference due to valid convolutions in the U-Net). From those predictions we extract synaptic partners on the fly with a *θ*_CC_ ≥ 0.95 (initial threshold of 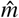, see 2.2) and store only their coordinates and associated connectivity scores if this score exceeds a very conservative threshold (*θ*_CS_ ≥ 5). To ensure sufficient context for partner extraction, we discard detections close (~100 nm) to the boundary of the prediction (≤28 voxels in xy, ≤2 sections in z). With a connected component radius of ~40 nm, this provides enough context to avoid duplicated connections at the boundary. A final block output size of (810, 810, 42) voxels per block or (3.2, 3.2, 1.6) μm results in a total of ~1.8 million blocks to cover the whole neuropil.

#### 3.3.1 Parallel processing and runtime

Using Daisy^5^ we parallelized processing with 80 workers, each having access to one RTX 2080Ti GPU for continuous prediction and five CPUs for simultaneous, parallel data pre-fetching and synaptic partner extraction. Processing all ~1.8 million blocks took three days, with a raw throughput of 1.53 μm^3^ s per worker, yielding a total of 244 million putative connections (see Fig. 1).

#### 3.3.2 Mapping predicted synapses to neuron skeletons

Given a neuron segmentation *l* : Ω ↦ *N*, a straight-forward mapping results from assigning each synaptic site to the neuron skeleton that shares the same segment (Fig. 5). However, false merges in the segmentation and misplaced skeleton nodes can lead to ambiguous situations where the segment containing a synaptic site contains several skeletons. Therefore, we assign synaptic sites to the closest skeleton node in the same segment, limited to nodes that are within a distance of 2 μm. The latter threshold was found empirically to confine the influence of wrong merges in the segmentation that otherwise could spread into neurites without skeleton traces. Wrong splits in the segmentation can lead to situations where no neuron skeleton shares the same segment as a synaptic site. In these cases, we leave the synaptic site unmapped.

**Figure 5:**
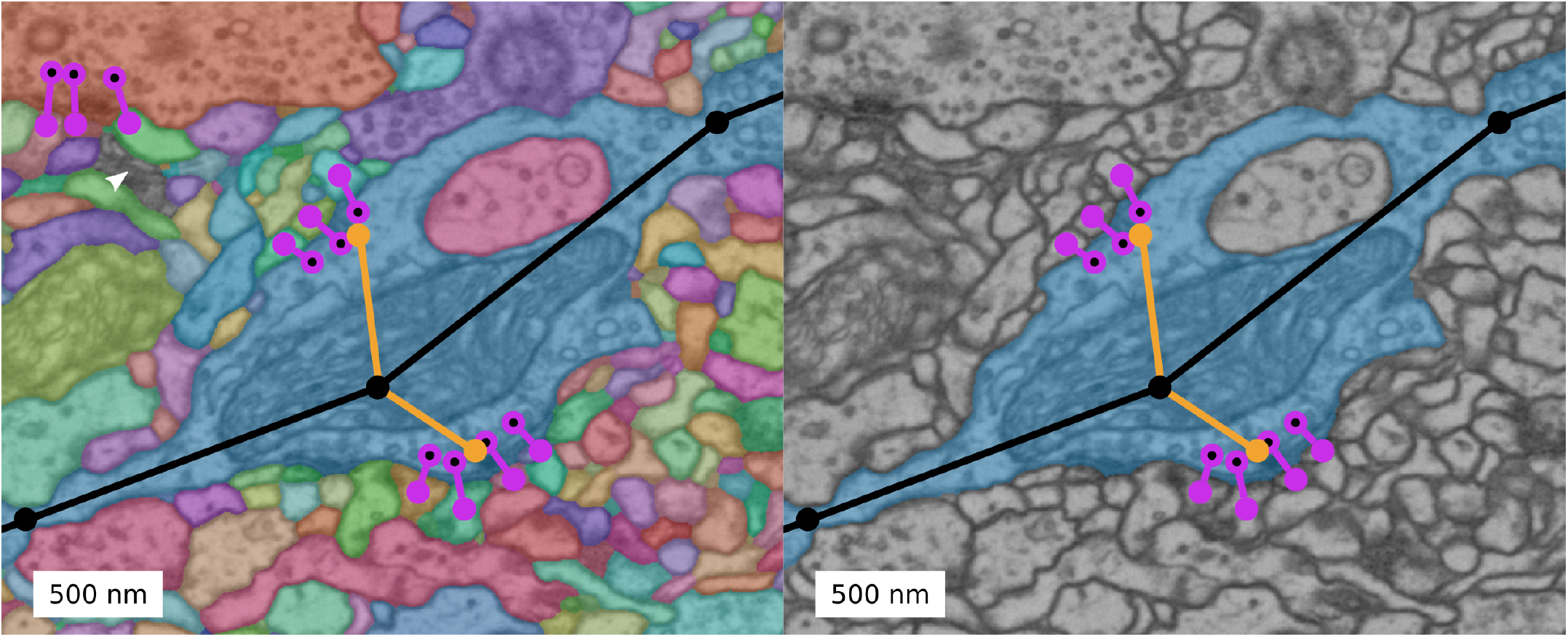
Mapping of predicted connections to manually created skeletons using neuron segmentation. Left: Around a manually traced skeleton (black line with nodes), we generate a neuron segmentation. Right: Predicted synaptic sites that intersect with the same neuron segment as the skeleton are assigned to that skeleton. For evaluation, we also use manually traced pre- and post-synaptic sites (orange nodes) for mapping. White arrowhead indicates zero-background representing putative thin glial processes.

According to the validation results on the Cremi dataset (see Section 3.1), we further filter connections inside a neuron. While for the Cremi dataset, we filter connections inside the same segment 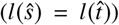, we here filter connections mapped to the same neuron 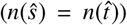. With this, we remove putative false positives, at the expense of potentially losing the ability to detect autapses (synapses connecting the identical neuron). We further cluster synaptic partners that connect the same neurons within a distance threshold of 250 nm, in order to avoid counting a single synaptic connection between two neurons multiple times. We remove all connections except the one with the highest score (*θ*_CS_) in each identified cluster.

### 3.4 Evaluation

#### 3.4.1 Datasets

We evaluate the quality of the whole-brain predictions on six comparatively large datasets with manually placed skeleton and synapse annotations annotated in Catmaid (Schneider-Mizell et al., 2016). We refer to these datasets in the following as InCalyx, OutLH, InOutEB, InOutPB, PairCalyx and PairLH. These datasets originate from four different brain regions: the calyx, the lateral horn (LH), the ellipsoid body (EB), and the protocerebral bridge (PB); see Fig. 4b for a rendering of their location within the full brain. Among those datasets, we distinguish two kinds of annotations: The first four datasets (InCalyx, OutLH, InOutEB, and InOutPB) contain manual skeleton traces of neurons that have all their incoming and/or outgoing synaptic connections annotated, possibly restricted to a well defined region (as reflected in the dataset name, *e.g.*, InCalyx refers to skeletons with all incoming synaptic connections annotated within the calyx). We refer to this kind of dataset as *synapse complete*. The remaining two datasets (PairCalyx, PairLH) consists of skeleton traces and synapse annotations of two sets of neurons, with known number of synaptic connections between every pair of neurons from one set to the other. We refer to this kind of dataset as *pair complete*. Statistics about all six datasets are shown in Table 1. These datasets include a diverse set of neuron types (see examples in Fig. 6b) and synaptic connections exhibiting different characteristics (see examples in Fig. 4: (c, d) shows polyadic axodendritic connections; (e) shows a single axo-axonic connection).

**Table 1:** Ground-truth datasets in FAFB. Cremi dataset was used for training (Section 2.4) and validation (Section 3.1), other datasets were used for evaluation (Section 3.4). Completeness column describes the nature of ground-truth annotation: **dense** means that an entire volume is densely annotated; **input** and/or **output** refer to *synapse complete* neurons for which all input and/or output connections are annotated; **connectivity** describes *pair complete* neurons for which all connections between two sets of neurons are annotated.

| Dataset | Completeness | Connection count | Neuron count (length [mm]) | Brain Region | Source |
| --- | --- | --- | --- | --- | --- |
| CREMI | dense | 1,965 | - | calyx | https://cremi.org |
| INCALYX | input | 59,155 | 528 (213) | calyx | Zheng et al. (2019), *in preparation* |
| OUTLH | output | 11,429 | 3 (2) | lateral horn | Bates et al. (2020) |
| INOUTEB | input & output | 61,280 | 27 (38) | ellipsoid body | Turner-Evans et al. (2019) |
| INOUTPB | input & output | 14,779 | 27 (16) | protocerebral bridge | Turner-Evans et al. (2019) |
| PAIRCALYX | connectivity | 43,595 | 144 (43), 528 (213) | calyx | Zheng et al. (2018) |
| PAIRLH | connectivity | 24,846 | 389 (243), 389 (243) | lateral horn | Bates et al. (2020) |

**Figure 6:**
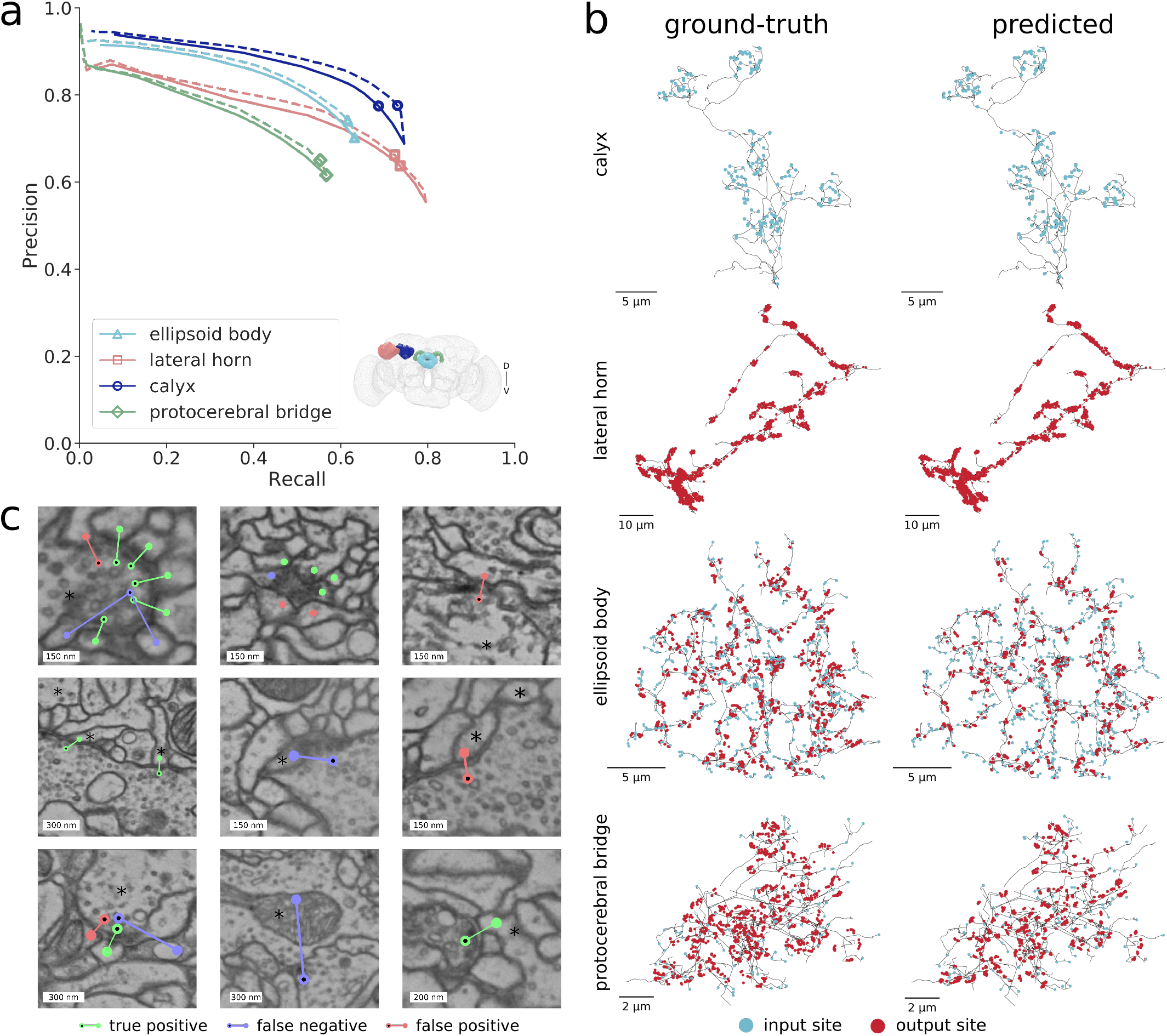
Evaluation on synapse-complete neurons in whole-brain dataset. **a**) Precision-recall curves for calyx (InCalyx), lateral horn (OutLH), ellipsoid body (InOutEB), protocerebral bridge (InOutPB). Adding synaptic cleft predictions as score (dashed lines) improves performance with original best f-score: 0.73, 0.68, 0.66, 0.59 versus f-score with synaptic clefts: 0.75, 0.69, 0.67, 0.60 **b**) Example neuron for each of the four datasets with ground-truth (left column) and predicted (right column) synaptic connections. We only display neuron parts and connections used for evaluation *i.e.* incoming connections in first row and outgoing connections in second row are omitted and neurons are intersected with respective brain regions. Overall, distribution of predicted synaptic sites along the neuron skeleton matches the ground-truth distribution. **c**) Examples of identified true positives, false positives, false negatives in lateral horn (top row), calyx (middle row), PB (bottom row) evaluated neuron marked with an asterisk. False positive in top row left, false positives and false negative in top row middle are examples of ambiguous cases, false negative in bottom row middle is an example of a missed axo-axonic connection.

InCalyx contains 528 Kenyon cells for which inside the calyx all 59,155 input connections have been annotated. A second dataset PairCalyx contains the same 528 Kenyon cells receiving input from 144 olfactory projection neurons (43,595 connections). The projection neurons are published by Zheng et al. (2018), Kenyon cells are unpublished (Zheng et al. (2019), in preparation).

OutLH contains three olfactory projection neurons, for which all 11,429 outgoing connections have been annotated inside the lateral horn. Synapses appear similar to PairCalyx, as both datasets contain the same type of neurons (compare Fig. 4c and d). A second dataset PairLH contains 389 neurons (including the three projection neurons from OutLH) with complete known connectivity inside the lateral horn (24,846 connections). The three olfactory projection neurons are published by Huoviala et al. (2018), remaining data is published by Bates et al. (2020).

InOutEB contains 27 neurons, for which all 61,280 incoming and outgoing connections have been annotated inside the EB. Data is published by Turner-Evans et al. (2019).

InOutPB contains the same 27 neurons as in InOutEB but neuron parts with all their 14,779 incoming and outgoing connections are located in PB. Data is published by Turner-Evans et al. (2019).

#### 3.4.2 Evaluation procedure

We evaluate the accuracy of predicted synaptic partners in two different ways, corresponding to the kind of ground-truth annotations we have available: For *synapse-complete* neurons, *i.e.*, neurons that (possibly only within a well defined region) have all their incoming and/or outgoing synaptic partners annotated, we evaluate the precision and recall of all synaptic partners that were mapped to those neurons. This evaluation applies to datasets InCalyx, OutLH, InOutEB, and InOutPB. For pairs of neurons with known connectivity, we evaluate the accuracy of correctly predicting the number of synaptic sites between those neurons (datasets PairCalyx and PairLH).

For both evaluation types, we first map each predicted pre- and post-synaptic site 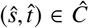 to a skeleton *n* ∈ *N* as described in Section 3.3. We will write 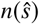 and 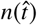 to refer to the mapped skeleton of 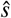 and 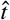, respectively. To guide the mapping, we generate a neuron segmentation locally around the synaptic sites to be matched (processed in 2 μm blocks with additional context of 1 μm to avoid border artifacts), using hierarchical agglomeration (Funke et al., 2018) favoring oversegmentations on affinity predictions obtained from Local Shape Descriptors (Sheridan et al., 2019). We also find it necessary to include manually traced pre-and post-synaptic sites for mapping. This is essential for partnering neurons for which no neuron skeletons exist. In this case, predicted connections are mapped onto single synaptic site nodes. We exclude connections from downstream analysis when either pre- or post-synaptic site connects to a putative glial cell (zero-valued background in the neuron segmentation). See Fig. 5 for an example segmentation and mapping, zero-background marked with a white arrowhead.

To evaluate precision and recall on synapse-complete neurons, we use the Cremi evaluation procedure (with skeletons IDs instead of neuron segment IDs) with the following two modifications to account for differences in synapse annotation: First, we increase the matching threshold for pre-synaptic sites from 400 nm to 700 nm. This more permissive threshold was empirically found to be necessary to compensate for the larger variance of pre-synaptic site placement in the one-to-many annotations used in the ground-truth (see Fig. 4c-f for examples). Second, we do not require a predicted post-synaptic annotation to be within a certain threshold distance to a post-synaptic site in the ground-truth, since the ground-truth annotations do not make use of a dedicated post-synaptic site marker. Instead, pre-synaptic nodes are directly connected to a skeleton node of the post-synaptic neuron, which is potentially far away from the predicted post-synaptic site. In summary, we consider a match between 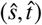 and a ground-truth annotation (*s*, *t*) possible, if 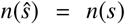, 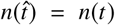, and 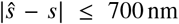. As in the original Cremi evaluation procedure, we perform a Hungarian matching to find at most one-to-one correspondences between possible matches, minimizing their Euclidean distance.

To measure the accuracy of correctly predicting the number of synaptic connections between pairs of neurons, we directly use the result of mapping predicted synaptic sites to skeletons. Let *w*(*n*_1_, *n*_2_) be the true number of synaptic partners between neurons *n*_1_ and *n*_2_ and

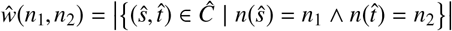

the number of predicted synaptic partners. We will refer to *w* and 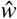 as the true and predicted *weight* of an edge between neurons. Given a weight threshold *γ*, we report the accuracy for correctly predicting an edge with at least that weight. However, most neuron pairs in the test set do not have a synaptic connection, which would lead to an unreasonably high accuracy stemming from trivially predictable negatives. Therefore, we limit the accuracy analysis to *relevant* neuron pairs {(*n*_1_, *n*_2_) | *w*(*n*_1_, *n*_2_) > 0}, but nevertheless count each non-relevant pair (*n*_1_, *n*_2_) for which 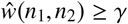 as an additional false positive.

We compare our method to a neuron-proximity baseline using neuron contact area as a proxy for synaptic weight. We compute this baseline by randomly generating synaptic connections. Specifically, we randomly select points using them as post-synaptic sites, and we obtain corresponding pre-synaptic sites by randomly drawing direction vectors from the predicted synaptic partners in FAFB. We then follow the same methodology as for the predicted connections with regards to synapse mapping (Section 3.3.2), and evaluation described above. As this procedure includes the removal of connections with pre- and post-synaptic sites mapped to the same neuron, we thus consider only those connections that link different neurons.

#### 3.4.3 Evaluation on Synapse-Complete Neurons

We evaluate our method on a fairly large dataset of synapse-complete neurons with a total of 146,643 synaptic connections (for comparison, the training and validation dataset contained 1402 and 540 synaptic connections, respectively). Precision-recall curves over the connection score threshold *θ*_CS_ are shown in Fig. 6a. Best achieved f-scores are **0.73** for InCalyx, **0.68** for OutLH, **0.66** for InOutEB and **0.59** for InOutPB. We observe highest accuracy for dataset InCalyx, which is proximal to the Cremi datasets we used for training and validation. In fact, the f-score of 0.73 for InCalyx is closest to the result on the validation set (f-score 0.76).

We further test whether the proposed method of scoring connections benefits from an independent synaptic cleft prediction. To this end, we use the synaptic cleft predictions from Heinrich et al. (2018) to derive a *cleft score* for each putative pair of synaptic partners by taking the maximal synaptic cleft value along a line from the pre- to the post-synaptic site. We then use the product of the connection score (see Section 2.2) and the cleft score to score and threshold synaptic partners. Results are shown in Fig. 6a as dashed lines. We observe a modest, but consistent, improvement for all datasets: **0.75** (+0.02) for InCalyx, **0.70** (+0.01) for OutLH, **0.67** (+0.01) for InOutEB and **0.60** (+0.01) for InOutPB.

#### 3.4.4 Evaluation on neuron connectivity

We make use of the ground-truth datasets PairCalyx and PairLH to evaluate our method in the context of automatically inferring a connectome.

The number of detected synapses depends on the score threshold *θ*_CS_. In order to obtain *θ*_CS_, we split neurons in PairCalyx and PairLH into a validation and test set. We use *θ*_CS_ that optimizes the f-score on the validation set and only use neurons in the test set for connectivity analysis. For PairCalyx, we use 105 Kenyon cells as validation set obtaining *θ*_CS_ = 60 (10 for synaptic cleft score). Our connectivity test set consists of 144 Projection neurons partnering with the remaining 423 Kenyon cells. For PairLH we use all three projection neurons in OutLH and obtain *θ*_CS_ = 60 (30 for synaptic cleft score). Our test set consists of the remaining 386 × 386 neurons.

We show connectivity results in terms of edge accuracy for different edge thresholds *γ* in Fig. 7d (using the metric described above in this section). At a reasonable threshold of *γ* = 5 we find an edge accuracy for PairCalyx of **0.96**, *i.e.* of the 2,279 ground-truth edges (2,039 pairs of neurons having five or more synaptic connections: *w* ≥ 5, 240 having less: *w* < 5) we find 1,965 to be correctly predicted as positive edge (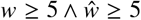, true positive) and 212 correctly predicted as having no edge (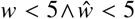, true negative). 28 pairs of neurons (one percent) have been incorrectly predicted as having a positive edge (false positive) and we find 74 pairs of neurons (three percent) to be incorrectly predicted as having no edge (false negative, missing edge in the connectome).

**Figure 7:**
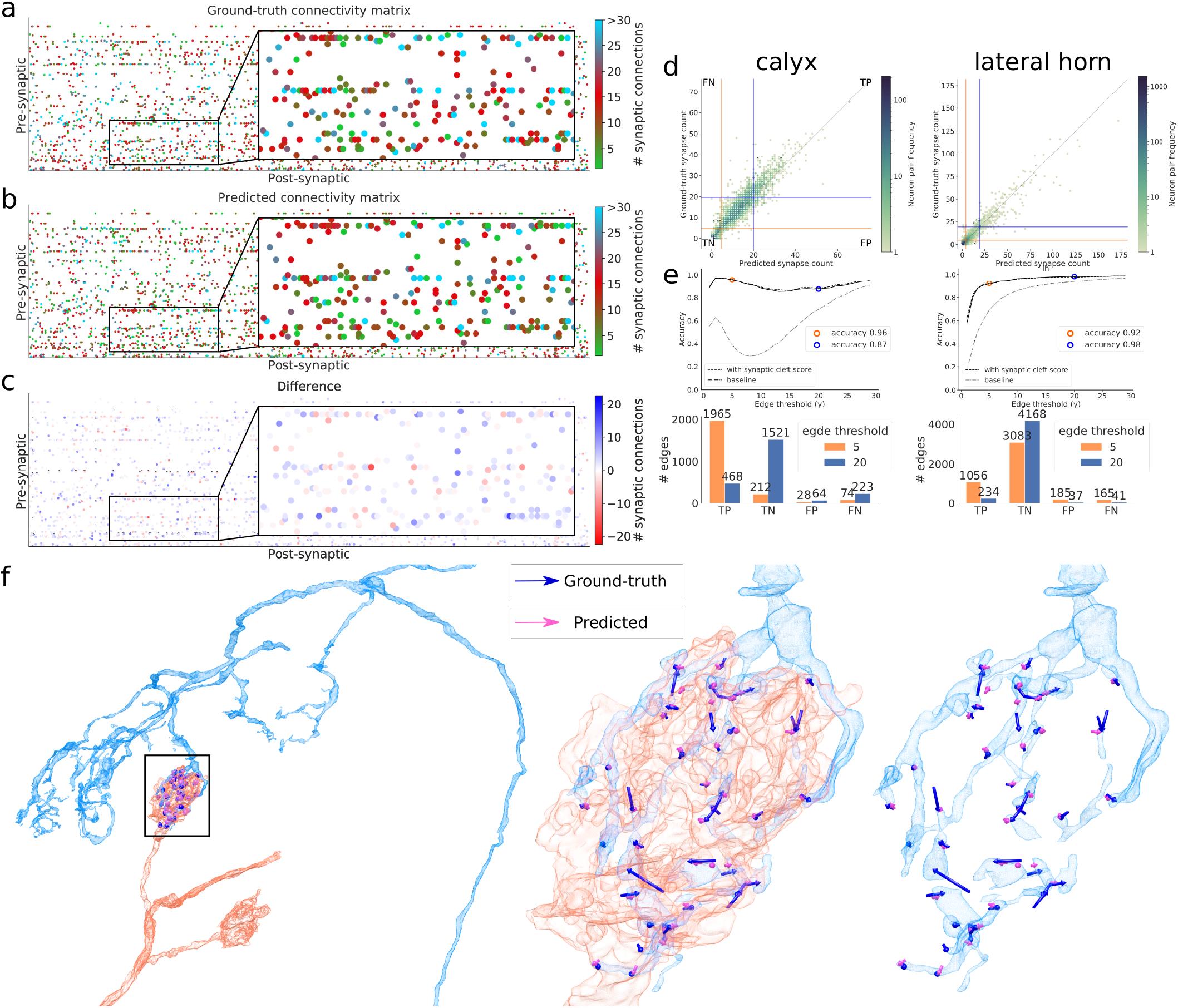
Comparison of automatically inferred versus ground-truth connectome. **a, b, c**) Ground-truth, predicted, and difference of number of synaptic connections between 144 projection neurons (pre-synaptic) and 528 Kenyon cells (post-synaptic) in calyx. **d**) For calyx, same data as in (a) and (b), shown as ground-truth versus predicted count of number of synaptic connections between pairs of neurons (for lateral horn, corresponding connectivity matrix is not shown). Pairs with both zero connections in ground-truth and prediction are omitted. Orange/blue line quadrants contain false negative (FN), true positive (TP), false positive (FP), and true negative (TN) edges for a thresholded connectivity criterion, *i.e.*, only edges with a synapse count equal or greater than *γ* = 5 (orange) or *γ* = 20 (blue) are considered true connections. **e**) Accuracy of thresholded connectivity criterion over different thresholds *γ* for calyx (left) and lateral horn (right). Results for thresholds *γ* = 5 and *γ* = 20 are highlighted and their absolute numbers of TPs, TNs, FPs, FNs is provided in the bar plots below. For comparison, we also show edge accuracy curves for the neuron-proximity baseline (see Section 3.4.2). **f**) Qualitative example of one edge in the connectome between a projection neuron (red) and a Kenyon cell (blue) with a ground-truth synapse count of 41 (blue arrows) and a predicted of 35 (pink arrows).

For PairLH, we obtain an edge accuracy of **0.92** for an edge threshold of *γ* = 5. We find that for both datasets the neuron-proximity baseline has a lower edge accuracy score of **0.39** (−0.57) for PairCalyx and of **0.69** (−0.23) for PairLH. Qualitatively, we see that the overall appearance of the predicted connectivity matrix for PairCalyx matches the appearance of the ground-truth connectivity matrix (Fig. 7a).

### 3.5 Data Dissemination

We implemented the Catmaid extension CircuitMap^6^ to make all predicted synaptic partners easily accessible in current circuit mapping workflows. Starting from a skeleton of interest, a Catmaid user can automatically retrieve all pre- and post-synaptic synaptic partners and link them with the selected skeleton. Additionally, and in conjunction with the availability of automatically generated skeletons from a neuron segmentation (Li et al., 2019), all up- and down-stream partner skeletons can be automatically imported. The CircuitMap extension will be deployed in Catmaid instances for the FAFB reconstruction community.

Furthermore, we provide an example Jupyter notebook together with the raw synaptic partner data for flexible, interactive analyses. The data also includes scores and segment identifiers, which were inferred for both pre- and post-synaptic locations based on the recent whole-brain neuron segmentation datset by Li et al. (2019). The notebook is part of the FafbSeg-Py^7^ package, and shows examples of querying the supervoxel graph based on synaptic connectivity or retrieval of synaptic connections for a given set of skeletons.

All predicted connections can also be directly explored in the browser-based volume viewer Neuroglancer^8^ by following www.tinyurl.com/tdq6xkw.

## 4 Discussion

### 4.1 Prediction Accuracy in FAFB

The results presented in the previous section indicate that the proposed method reliably detects connectome edges with five or more synaptic connections outperforming the neuron-proximity baseline by a large margin (edge accuracy: 0.96 versus 0.39 for PairCalyx and 0.92 versus 0.69 for PairLH, see Section 3.4). Qualitatively, we find those results confirmed across different neuron types from all four evaluated datasets: As shown in Fig. 6b, the overall distribution of synaptic sites along the skeleton is generally preserved. In particular, the distribution of predicted pre- and post-synaptic sites of more complex neuron morphologies in the ellipsoid body and the protocerebral bridge agree with the ground-truth distribution.

The high accuracy of detecting connectome edges in dataset PairCalyx and qualitative impressions seem to contrast the comparatively low f-scores obtained on the synapse-complete dataset InCalyx on the same brain region (0.73 without using a cleft prediction and 0.75 with a cleft prediction). For comparison, the example neuron of the calyx shown in Fig. 6b (top row) has a per-neuron f-score of 0.79, despite the fact that prediction and ground-truth largely agree.

We attribute this discrepancy largely to the fact that the Cremi metric we use to evaluate the synapse-complete datasets is quite conservative (for comparison, also the current leader of the Cremi challenge has a comparatively low f-score of 0.58): due to the Hungarian matching performed between true and predicted synaptic pairs, a predicted connection where at least one of the synaptic sites is slightly incorrectly placed such that it ends up on a different neuron segment adds both to the number of false positives and false negatives. The edge accuracy, on the other hand, would at most count one missing connection (which might even be compensated for by an additional detection further away). As such, the Cremi metric is more sensitive to errors in the neuron segmentation (especially in proximity to synaptic clefts) and spurious or missing manual annotations in ambiguous situations (see Fig. 6c, first row middle for questionable false positives and false negatives). Furthermore, false merges in the neuron segmentation cause correctly predicted synaptic connections to be incorrectly mapped onto a skeleton, and thus contribute to the number of false positives of the evaluated skeleton. This type of error will be counted in the Cremi metric for each neuron, but in the edge accuracy only if the merger occurs between a pair of neurons contained in the evaluation dataset. Similarly, since the edge accuracy is limited to pairs of neurons with known connectivity, false positives to other neurons are not counted either.

Hence, the Cremi metric should be interpreted as a measure of the whole pipeline accuracy, *i.e.*, not just of the prediction of synaptic partners, but also of the neuron segmentation, of the exact placement of skeleton nodes, and of the mapping of synaptic partners to skeletons. Given a perfect neuron segmentation, the true precision and recall values of the synaptic partner prediction alone would therefore be higher than reported.

Consequently, the overall accuracy of the proposed method will benefit from more accurate neuron segmentations, in particular within the proximity of synaptic sites. Since the prediction of synaptic partners does not require a neuron segmentation (a segmentation is only needed for the mapping of synaptic sites to skeletons), the segmentation can be replaced with a more accurate one in the future to improve the mapping, without having to retrain or reprocess the synaptic partner predictions.

Despite the fact that the absolute value of the Cremi metric is a lower bound to the actual synapse prediction accuracy, we observe a significant decrease in f-score in brain regions farther away from the calyx. This effect is most notable in terms of increased false negatives in the protocerebral bridge (InOutPB f-score: 0.59), which anecdotally stem from failures to detect axo-axonic links (via manual inspection, see Fig. 6c bottom row middle for an example). Less pronounced but still noticeable, we also observe a decrease in performance when comparing the edge accuracy obtained in calyx compared to lateral horn (0.96 versus 0.92). Given that all the training data used was cropped from the calyx, the decrease in performance suggests that the phenotype of synapses is not uniform across neuron types and that the network learnt to recognize neuron type-specific features.

Our observations have two implications for future work: First, training and validation should be carried out on more diverse datasets that capture more of the variation of synapse phenotypes throughout the whole brain. In particular, connectivity accuracy should be manually validated for regions and neuron types that were not part of the training dataset and error estimates should be incorporated into downstream analysis. Second, the quality of a neuron segmentation should be evaluated not just based on overall topological correctness, but also on accuracy close to boundaries, in particular in the proximity of synaptic clefts. Neither of the two most commonly used metrics to evaluate neuron segmentations (expected run length and variation of information) are sensitive to small errors close to synaptic terminals (Plaza and Funke, 2018).

### 4.2 Model validation on Cremi

#### 4.2.1 Multi-Task networks achieve same performance as single-task networks

We show that the multi-task network MT_2_ (two separate upsampling paths for 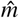 and 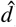) achieves similar performance compared to two having separate networks (ST), despite having 40 % less parameters. This is somewhat surprising, since the baseline multi-task network MT_1_ (single U-Net with two outputs for 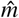 and 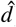) failed to converge either for 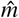 or 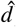. A similar observation has been made by Wang et al. (2019), who independently to our work compared models similar to our ST, MT_1_ and MT_2_ on a different computer vision task and also found MT_2_ (called Y-Net in their work) to perform best. Further improvements in training multi-task networks are possible by exploring different weighting schemes (Kendall et al., 2018; Wang et al., 2019) other than simply summing or multiplying both losses as we did here. Already, MT_2_ offers a promising starting point for a general volumetric EM U-Net, where multiple different tasks are jointly solved (*e.g.*, the segmentation of boundaries, intracellular organelles, synapses).

#### 4.2.2 Impact of mini-batch sampling during training

Due to the sparsity of synaptic sites, we found that rejection of mini-batches that do not contain synapses is important to prevent the neural networks from converging to trivial solutions (*i.e.*, predicting zero for the post-synaptic site mask). If we train with constant *p*_rej_ = 0 (no mini-batches are rejected during the entire training), success or failure highly depends on the strength of voxel-wise balancing (data not shown). We generally observed stable training when we reject empty mini-batches with a probability of *p*_rej_ = 0.95 (as described in Section 2.4). Furthermore, we found overall better validation performance when using a “curriculum” strategy, *i.e.*, we first train with a rejection probability of *p*_rej_ = 0.95 until 90k iterations and *p*_rej_ = 0 afterwards. The average f-score of all 16 tested setups for a constant rejection probability is 0.71±0.02 compared to 0.73±0.02 for the curriculum strategy (see f-score results in top two rows and bottom two rows in Fig. 3). A likely explanation for the increase in accuracy is that with a constant rejection probability of *p*_rej_ = 0.95, mini-batches containing synapses are overrepresented during training and the resulting networks are weaker in correctly classifying negatives in areas that do not contain synapses such as the cell interior of large neurons.

### 4.3 Limitations

Our model is well-suited to detect one-to-many synapses (one pre-synaptic site targeting many post-synaptic sites), as it explicitly extracts an individual connection for each detected post-synaptic site. It is however less suited to detect the relatively rare many-to-one synapses, which can be found for instance in the α-lobe of the mushroom body (Takemura et al., 2017). For instance, if we use connected components to find post-synaptic sites in 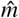, we are likely to only find a single site for a many-to-one synapse and thus would also only extract a single connection. To capture more complex synapse configurations will be the subject of future work.

## Acknowledgements

We thank Scott Lauritzen for helping with data acquisition; Matthew Nichols for importing Catmaid annotations; William Patton for code contributions; Nils Eckstein for helpful discussions; Jeremy Maitin-Shepard for adding synapse visualization features to neuroglancer; Vivek Jayaraman for providing evaluation data. We would also like to thank Zhihao Zheng, Feng Li, Corey Fisher, Nadiya Sharifi, Steven Calle-Schuler and Davi Bock for access to prepublication data used for evaluation.

## Author contributions

*Conceptualization*: Srinivas Turaga, Jan Funke. *Funding acquisition*: Wei-Chung Allen Lee, Rachel Wilson, Matthew Cook, Jan Funke. *Software*: Julia Buhmann, Arlo Sheridan, Stephan Gerhard, Renate Krause, Tri Nguyen, Larissa Heinrich, Philipp Schlegel, Stephan Saalfeld, Jan Funke. *Validation and evaluation*: Julia Buhmann, Jan Funke. *Evaluation data generation*: Philipp Schlegel, Gregory Jefferis, Davi Bock. *Data dissemination*: Julia Buhmann, Stephan Gerhard, Jan Funke. *Supervision*: Matthew Cook, Jan Funke. *Visualization*: Julia Buhmann, Arlo Sheridan, Jan Funke. *Writing original draft*: Julia Buhmann, Jan Funke. *Writing - review & editing*: Julia Buhmann, Arlo Sheridan, Stephan Gerhard, Renate Krause, Tri Nguyen, Larissa Heinrich, Philipp Schlegel, Stephan Saalfeld, Gregory Jefferis, Davi Bock, Srinivas Turaga, Matthew Cook, Jan Funke.

## Footnotes

1. https://github.com/funkelab/synful_fafb

2. https://github.com/flyconnectome/fafbseg-py

3. https://github.com/funkey/gunpowder

4. https://cremi.org

5. https://github.com/funkelab/daisy

6. https://github.com/BrainCircuitsIO/circuitmap

7. https://github.com/flyconnectome/fafbseg-py

8. https://github.com/google/neuroglancer

